## Supplemental material for "HYPOTHESIS GENERATION FOR RARE AND UNDIAGNOSED DISEASES THROUGH CLUSTERING AND CLASSIFYING TIME-VERSIONED BIOLOGICAL ONTOLOGIES"

### 1. Supplemental Network Clustering

Considering the generalizability of networks, many fields have devoted efforts to modeling network structure including statistics, physics, computer science, biology, and social sciences. Within biology, networks have been used successfully for well over a decade now. The applications have been far and wide including disease variant prioritization[1;2], drug-target identification[3;4], disease gene detection[5]. One sub-field of network science commonly used within a biological context is network clustering.

Network clustering is the general process of assigning nodes to collections whose members are densely connected within the collection yet sparsely connected to the rest of the network. These relatively dense subsets of nodes are called clusters (or communities)[6]. The utility of network clustering was arguably established when cluster members were shown to embody structural and functional similarities[7]. As a result, a flurry of algorithms designed to partition a graph to create this disparity in connectedness within and between partitions was proposed[8;9].

The rapid growth of literature prompted several comparative studies in the late 2000s[10;11] and generalizations in the 2010s[12;13;14]. While the literature is expansive, common approaches to network clustering include those algorithms 1) based on topological distance or random walks, 2) based on the optimization of some objective function such as modularity or surprise, and 3) based on statistical models or principles. While there are many available options for

[1]Department of Computer Science, University of Colorado Boulder, Boulder, CO, USA 80309,

[2]Department of Statistics, Colorado State University, Fort Collins, CO, USA 80523,

[3]Precision Medicine Institute, Children's Hospital Colorado, Aurora, CO, USA 80045,

[4]Department of Biostatistics & Informatics, Colorado School of Public Health, Aurora, CO, USA 80045,

*Date*: November 9, 2023.

network clustering, the algorithms chosen for this study cover each of the three general approaches and have been shown to outperform competing methods in comparative studies.

We focus on methods designed to uncover disjoint and overlapping clusters (e.g., see Figure 1A and B). The disjoint clustering methods considered include a greedy modularity maximization method [15], walktrap [16], and infomap [17;18]. While each method seeks to partition the node set, the objective differs across each method, leading to fundamentally different clusters. CESNA (communities from edge structure and node attributes) [14] is an overlapping clustering method that leverages both the topology of the network and node metadata to form clusters. Unlike the greedy, walktrap, and infomap methods, CESNA distinguishes between genes and phenotypes, treating the biological network as a heterogeneous graph. A summary of each method is included in the following subsections.

**Greedy.** Proposed in [15], this clustering method seeks to optimize a modularity score. In particular, a partition creating the largest disparity between the fraction of edges internal to the partition and the expected fraction from a random graph with the same degree sequence is desired. Since maximizing this quantity is NP-complete in the strong sense [19], a greedy heuristic is used to find reasonable clusters. To start, each node is considered a community. At each iteration of the algorithm, two communities that contribute maximum positive value to the global modularity score are merged. This process continues until no such increase in modularity is possible. The estimated complexity of this method on sparse networks is $\mathcal{O}(nlog^2n)$ where $n$ denotes the number of nodes in the network. Greedy is implemented in the Python networkx package [20].

**Walktrap.** Walktrap was introduced in [16] as a means for clustering a graph using random walks, positing that short random walks tend to remain within a community. To start, each node is considered a community. Distances between communities are computed via random walks, and communities are merged such that there are shorter walks within a community and larger walks between communities. This process is repeated $n - 1$ times implying a complexity of $\mathcal{O}(mn^2)$ where $m$ denotes the number of edges in the graph, or $\mathcal{O}(n^2logn)$ for sparse graphs [21]. Walktrap is implemented for Python in the CDlib library [22].

**Infomap.** Rooted in information theory, infomap discovers communities by minimizing the description length of an information flow on a graph, a variant of a coding problem [23]. Information flows are measured across a network using random walks where groups of nodes for which information flows easily "can be aggregated and described as a single well-connected module" [17]. The authors demonstrate these modules are synonymous with communities. The map equation, introduced in [18], provides the theoretical basis for the infomap algorithm, describing how well information

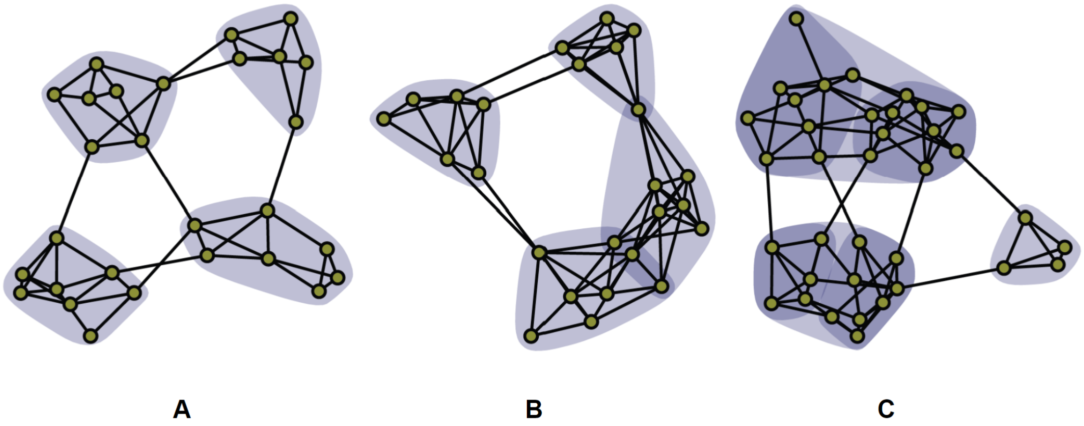


FIGURE 1. A contrived example illustrating various types of community structure including a) disjoint, b) overlapping, and c) hierarchical communities. Adapted from[27].

about the original network is transferred through a given network partition.[24] estimates that the complexity of infomap is $\mathcal{O}(m)$. This method is available through the infomap package in Python[25].

**CESNA.** Distinct from its competing methods, CESNA (communities from edge structure and node attributes) discovers communities using the structure of the edges within a graph and the available node attributes. In particular, CESNA treats the biological network as a heterogeneous network, distinguishing genes from phenotypes rather than treating them fundamentally the same. The method posits that a graph arises from its nodes' attributes and its nodes' community structure before inferring the latent community structure via maximum likelihood estimation under a statistical model[14]. Discovered communities should be topologically dense and maintain similar node attributes. One iteration of CESNA has an estimated complexity of $\mathcal{O}(m+nk)$ where $k$ is the number of node attributes. For this study, the node type (i.e. gene or phenotype) is the only node attribute considered implying a computational complexity of $\mathcal{O}(m + n)$. This method is implemented using the Stanford Network Analysis Platform (SNAP)[26].

### 2. Supplemental Network Subclustering

Clusters in real networks tend to exhibit hierarchical structure such that larger clusters are assumed composed of smaller clusters that can be further divided (e.g., see 1C). For this study, we leverage hierarchical clustering methods to subdivide clusters attained from the network clustering methods discussed in Section 1 into "subclusters" of manageable size using Paris[28]. This method produces a dendrogram which can be cut to produce subclusters. The dendrogram is cut such that the largest resulting subcluster maintains no more than 100 members. This subclustering method is described in the following subsections and implemented in the Python scikit-network package[29].

**Paris.** Paris is an agglomerative hierarchical clustering method based on node pair sampling. In particular, the method proposes a distance measure that compares the joint probability of sampling a pair of nodes, say $i$ and $j$, to the product of the marginal probabilities of sampling $i$ and $j$. In particular, two nodes are relatively close if the probability of sampling $j$ given that $i$ has been sampled is large compared to the probability of sampling $j$. To start, each node is assigned to its cluster. Recursively, the two closest clusters are combined resulting in a dendrogram. The algorithm is a modification of the Louvain algorithm where the iterative step is replaced by a single merge. The complexity of paris is $\mathcal{O}(m)$.

Each clustering and subclutering method is summarized in terms of its class, complexity, and objective in Table 1, along with relevant references.

TABLE 1. Description of clustering and subclustering methods considered in this study. The complexity and method objectives are provided where $n$ denotes the number of nodes, $m$ denotes the number of edges, and $k$ denotes the number of node attributes. References are provided for each method.

| Class | Method | Complexity | Objective | References |
|---|---|---|---|---|
| Clustering | Greedy | $\mathcal{O}(n \log^2 n)$ | Modularity | 15 |
| | Walktrap | $\mathcal{O}(n^2 \log n)$ | Distance based on random walks | 16;21 |
| | Infomap | $\mathcal{O}(m)$ | Map equation | 23;18;24 |
| | CESNA | $\mathcal{O}(m + nk)$ | Latent space likelihood | 14 |
| Subclustering | Paris | $\mathcal{O}(m)$ | Distance based on node pair sampling | 28;30 |

### 3. Nontrivial Clusters

When applying greedy, walktrap, infomap, and cesna to the 2021 biological network, each cluster attained is larger than For the purposes of the study, only small communities maintain

Applying greedy, paris, infomap and cesna to our network of the 2021 data resulted in very few clusters being identified, most of which were very large in size Figure 2. The cesna clusters are an exception to this generalization, all of which are ¡ 200 members in size. Our goal is to recommend a set of genes and HPO terms to clinicians and experimentalists for further investigation, providing such users with a list of several thousand items is not a useful product. After discussion with clinicians we frequently collaborate with, we determined that to be useful clusters would need to have fewer than 100 nodes. We later found articles[31] from a DREAM challenge which required that all clusters be in the range of 3-100 nodes because ”[clusters] with over 100 genes are typically less useful to gain specific biological insights”[32]. To meet this new requirement, we re-clustered all previously identified clusters using

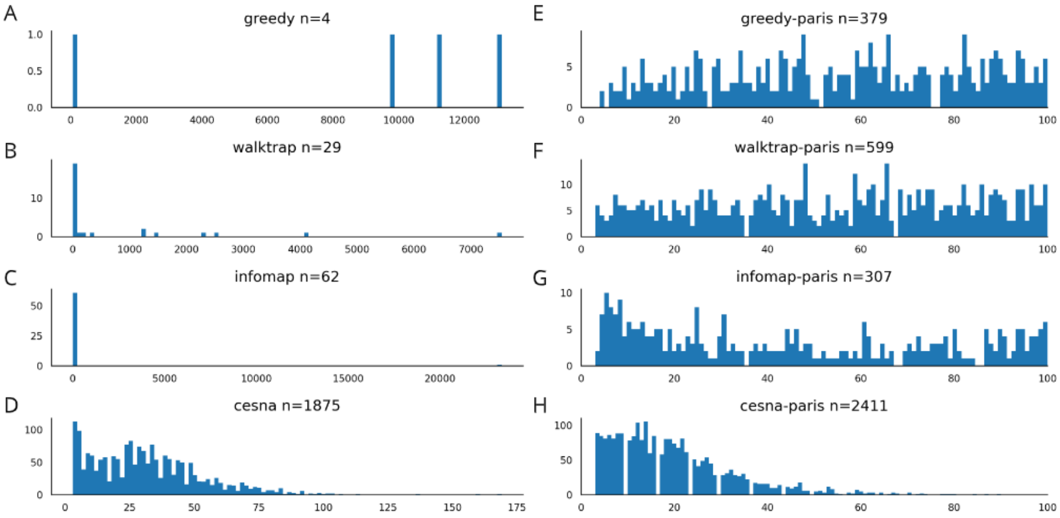


FIGURE 2. Shown are distributions of cluster size using A. greedy B. walktrap C. infomap and D. censa. The second column contains the subcluster size distributions, where every cluster from panels A-D were clustered again with the paris-hierarchical method E. greedy-paris F. walktrap-paris G. infomap-paris and H. censa-paris. In all of these plots the x-axis is cluster-size and the y-axis is the number of clusters. The first three of the clustering algorithms on their own, have a tendency to produce very few clusters that are all very large, some with a membership larger than 20,000 nodes, in the case of infomap. Our end goal is to use these clusters to provide sets of genes and phenotypes that are likely to have yet-to-be-discovered clinically meaningful relationships. These clusters in A-D are far too large to be useful for hypothesis generation in clinical or experimental settings. A second layer of cluster with the paris method is applied and shown in F-H, setting an upper limit of 100 on cluster size results in many more clusters of a size manageable for human curation.

Paris hierarchical clustering[28]. Paris allows an upper limit to cluster size to be set, this resulted in many more clusters than previously identified all $\leq 100$ nodes in size (Figure 2).

### 4. Data Sources

Two main data sources have been used in this study, namely STRING[33] and Human Phenotype Ontology (HPO)[34]. Deserving additional mention is the genes-to-phenotype annotation file produced by HPO which aggregates information from Online Mendelian Inheritance in Man (OMIM)[35] and Orphanet[36]. These data sources were gathered for the years 2019-2021. STRING was gather easily using their portal (https://string-db.org/cgi/access?sessionId=b8wDXZhU8UYD&footer_active_subpage=archive) for accessing older versions of the data, specifically only the Homo sapien (species code 9606) data was gathered and used. Collecting older versions of HPO was less straightforward. Older version of HPO were gathered from git commit histories of the Github repositories: obophenotype/human-phenotype-ontology and drseb/HPO-archive. The genes-to-phenotype annotation files were gathered from a mix of those same Github repositories, the Monarch Initiative Jenkins servers (current ones and an older one accessed via the Wayback Machine[37]). Specific URLs used for each of these data sources can be found in the reproducible pipeline Snakemake file[38]. The publication dates (or date ranges in the case of STRING) can be seen in Table 2. Due the unavailability of g2p annotations prior to 2019, we opt to use only data 2019 - 2022.

TABLE 2. The three data sources gathered and used in this study and the accompanying dates each version was originally published (or published and active in the case of STRING). All dates are in year-month-day format. Genes-to-phenotype 2019 is taken from the way-back machine. It cannot be automatically downloaded so I copy and pasted the information and saved it in a file in the git repository. Previous versions of STRING and well preserved and documented. The availability of previous versions of HPO and the G2P annotations do not have a central or documented location for previous versions, we use what is available and closest to the end of each calendar year. HPO and g2p date are time stamps associated with each files in the GitHub history. Of note is that genes-to-phenotypes 2019 is taken from the way-back machine. Automatically downloading it proved difficult so the information was copied, pasted, and saved in a file in the git repository. Additionally, genes-to-phenotypes 2018 could not be found in any GitHub repository histories or the way back machine, for reason we do not use data futher back than the end of 2019.

| Year | HPO | Genes-to-phenotype | STRING |
|---|---|---|---|
| 2019 | 2019-11-08 | 2019-9-2 | 2019-1-19 to 2020-10-17 |
| 2020 | 2020-12-07 | 2020-8-25 | 2020-10-17 to 2021-8-12 |
| 2021 | 2021-10-10 | 2021-10-10 | 2021-8-12 to present |
| 2022 | 2022-12-06 | 2022-12-06 | 2021-8-12 to present |

### 5. Null Model Comparison

When choosing the snowball sampling null model we compared it to two other null models, random clusters and edge shuffle, and found snowball sampling to be a far more rigorous model. We compared the of empirical p-values in the 2019 greedy-paris cluster p-values calculated using the three null models, see and their resulting distributions in Figure S3. With the random clusters null model 39% of clusters achieved significance ($p < 0.05$), and the edge shuffle null model resulted in 48%. Unlike these two other null models where a substantial portion of clusters were significant, we found snowball sampling to be very conservative in comparison, only 6% were significant and 70% of the clusters had $p = 1.00$.

The random cluster null model works by:

- Start with a real cluster, of size X.
- Choose X node uniformly at random from the network without replacement to create a synthetic cluster.
- Perform rediscovery on the synthetic cluster.
- repeat steps 2 and 3 1,000 times

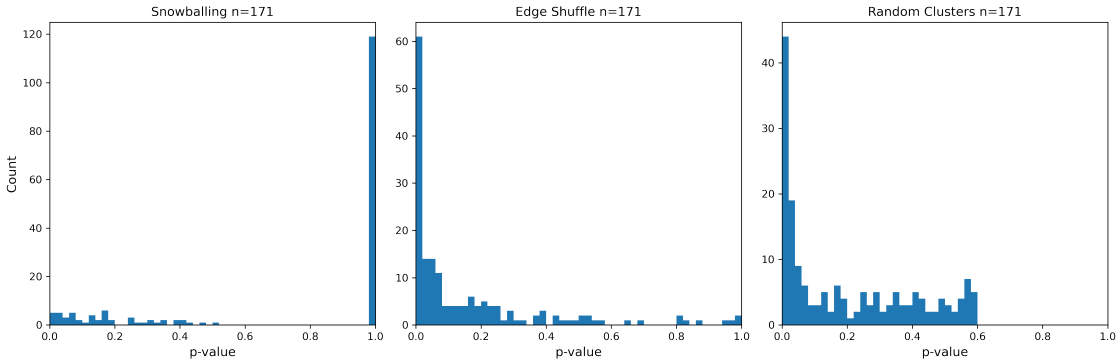


FIGURE 3. Distribution of empirical p-values of the 2019 greedy-paris clusters using three different null models: snowballing sampling, edge shuffle, and random clusters. All figures share the same y-axis which is also on a log scale. The proportion of clusters with $p < 0.05$ in each model is 6%, 48%, and 39%, for snowballing, edge-shuffle, and random clusters respectively. Snowballing has 70% of its clusters with $p = 1.00$, whereas the other two models have zero clusters falling into this category.

- Calculate the empirical p-value of the number of rediscoveries in the real cluster versus the number of rediscoveries in it's 1,000 synthetic clusters.

Edge shuffle null model tests the effect of randomized edge for rediscovery rather than randomized clusters. This model works by:

- For each real 2019 cluster
- Take the list of new edges added in 2020 and shuffle which nodes constitute a new synthetic edge - synthetic edges are comprised of the same nodes as the real new edges but in randomized pairings.
- Performed rediscovery on the real cluster with the list of synthetic edges.
- repeat steps 2 and 3 1,000 times
- Calculate the empirical p-value of the number of rediscoveries for each cluster with the real list of new edges versus the number of rediscoveries with 1,000 synthetic new edge list.
